# Partial Fitch Graphs: Characterization, Satisfiability and Complexity∗

**DOI:** 10.1101/2024.04.30.591842

**Authors:** Marc Hellmuth, Annachiara Korchmaros, José Antonio Ramírez-Rafael, Bruno Schmidt, Peter F. Stadler, Sandhya Thekkumpadan Puthiyaveedu

## Abstract

Horizontal gene transfer is an important contributor to evolution. Following Walter M. Fitch, two genes are xenologs if at least one HGT separates them. More formally, the directed Fitch graph has a set of genes as its vertices, and directed edges (*x, y*) for all pairs of genes *x* and *y* for which *y* has been horizontally transferred at least once since it diverged from the last common ancestor of *x* and *y*. Subgraphs of Fitch graphs can be inferred by comparative sequence analysis. In many cases, however, only partial knowledge about the “full” Fitch graph can be obtained. Here, we characterize Fitch-satisfiable graphs that can be extended to a biologically feasible “full” Fitch graph and derive a simple polynomial-time recognition algorithm. We then proceed to show that several versions of finding the Fitch graph with total maximum (confidence) edge-weights are NP-hard. In addition, we provide a greedy-heuristic for “optimally” recovering Fitch graphs from partial ones. Somewhat surprisingly, even if ∼ 80% of information of the underlying input Fitch-graph *G* is lost (i.e., the partial Fitch graph contains only ∼ 20% of the edges of *G*), it is possible to recover ∼ 90% of the original edges of *G* on average.

## 1 Introduction

Horizontal gene transfer (HGT) is a biological p rocess by which genes from sources other than the parents are transferred into an organism’s genome. HGT is an important contributor to evolutionary innovation in particular in microorganisms [5]. The identification of HGT events from genomic data, however, is still a difficult problem in computational biology [17], both in practical applications and in terms of the underlying mathematics and algorithmics. In most situations, it can be assumed that the evolution of genes is tree-like and thus can be described by a gene tree *T* whose leaves correspond to the present-day, observable genes; the interior vertices and edges then model evolutionary events such as gene duplications, speciations, and also HGT. Since HGT distinguishes between the gene copy that continues to be transmitted vertically, and the transferred copy, one can associate the transfer with the edge in *T* connecting the HGT event with its transferred offspring [9].

Recent formal results are phrased in terms of a binary relation defined between a pair of present-day genes [7]. The corresponding Fitch (xenology) graph contains an edge *x* → *y* whenever an HGT event occurred between *y* and the least common ancestor of *x* and *y* [7]. Fitch graphs form a hereditary sub-class of the directed cographs [4], and admit a simple characterization in terms of eight forbidden induced subgraphs on three vertices (see Figure 1 below) [7, 11]. Moreover, every Fitch graph uniquely determines a least resolved edge-labeled tree by which it is explained. This tree is related to the gene tree by a loss of resolution [7].

**Figure 1:**
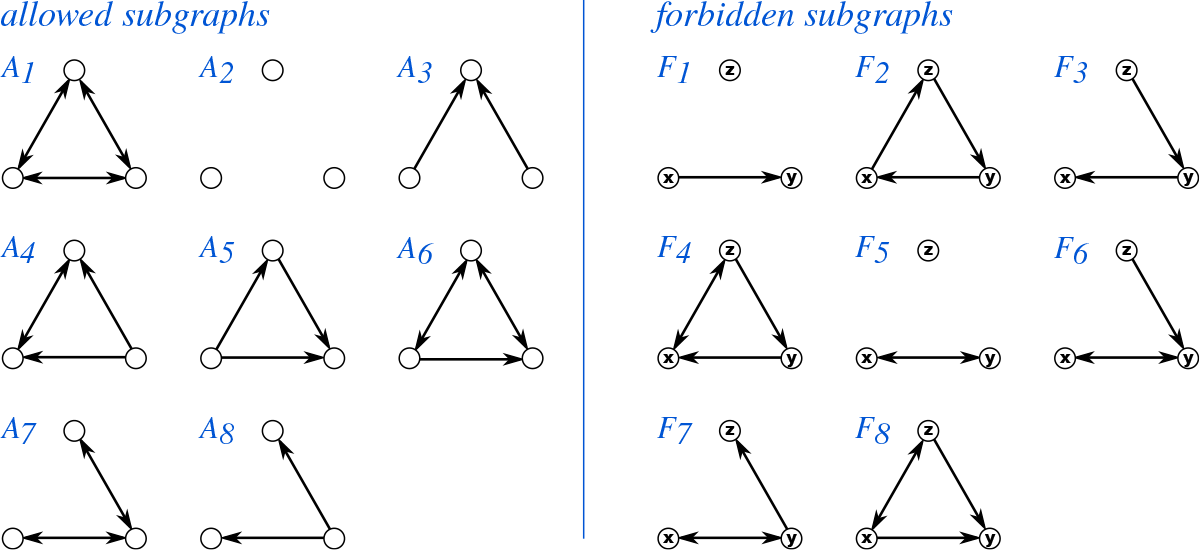
Fitch graphs are characterized by their induced subgraphs on three vertices: of the 16 possible irreflexive binary relations on three vertices, eight (*A*_1_ through *A*_8_) may appear in Fitch graphs, while the remaining eight (*F*_1_ through *F*_8_) form forbidden induced subgraphs.

Information on HGT events can be extracted from sequence information using a broad array of methods [17], none of which, however, is likely to yield a complete picture. Reliable information is needed to decide whether or not two genes are xenologs; thus, it may be available only for some pairs of genes (*x, y*), but not for others. In this situation, it is natural to ask whether such partial knowledge can be used to infer missing information. One can think about the latter problem as marking a subset *E*^*′*^ ⊆ *E* of edges in a digraph *G* = (*V, E*) with a “?”-symbol and then ask to what extent these “questionable edges” can be replaced by directed edges or non-edges to recover *E*. In [16], the analogous question was investigated for di-cographs. The main formal result of the present contribution is a characterization of partial Fitch graphs, Theorem 3.11. This characterization is then used to design an accompanying polynomial-time algorithm to decide whether or not a given graph *G* is a partial Fitch graph. In the affirmative case, a full Fitch graph *G*^*∗*^ can readily be obtained from the underlying trees that explain *G*. In addition, we show that the “weighted” version of Fitch graph completion is NP-hard, and we provide a simple greedy heuristic for finding “optimally” weighted Fitch graphs. We evaluate these algorithms on simulated data. As it turns out, only information about a very few but correct edges of the original Fitch graph *G* is needed to recover *G* fully.

## 2 Preliminaries

### Relations

Throughout, we consider only *irreflexive, binary* relations *R* on *V*, i.e., (*x, y*) ∈ *R* implies *x*≠ *y* for all *x, y* ∈ *V*. We write 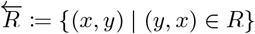 and 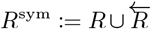 for the *transpose* and the *symmetric extension* of *R*, respectively. The relation 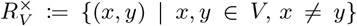 is called the *full* relation.

For a subset *W* ⊆ *V* and a relation *R*, we define the *induced* subrelation as *R*[*W*] := {(*x, y*) | (*x, y*) ∈ *R, x, y* ∈ *W* }. Moreover, we consider ordered tuples of relations ℛ = (*R*_1_, …, *R*_*n*_). Let 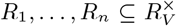, then ℛ = (*R*_1_, …, *R*_*n*_) is *full* if 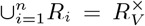 and *partial* if 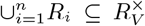. Note that a full tuple of relations is also considered to be a partial one. Moreover, we consider component-wise sub-relation and write ℛ[*W*] := (*R*_1_[*W*], …, *R*_*n*_[*W*]) and 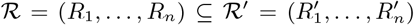 if 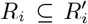 holds for all *i* ∈ {1, …, *n*}. In the latter case, we say that ℛ^*′*^ *extends* ℛ.

### Digraphs and DAGs

A directed graph (digraph) *G* = (*V, E*) comprises a vertex set *V* (*G*) = *V* and an irreflexive binary relation *E*(*G*) = *E* on *V* called the edge set of *G*. Given two disjoint digraphs *G* = (*V, E*) and *H* = (*W, F*), the digraphs *G*∪*H* = (*V* ∪*W, E*∪*F*), 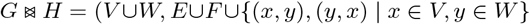) and *G ▷ H* = (*V* ∪ *W, E* ∪ *F* ∪ {(*x, y*) | *x* ∈ *V, y* ∈ *W* }) denote the *union, join* and *directed join* of *G* and *H*, respectively. For a given subset *W* ⊆ *V*, the *induced subgraph G*[*W*] = (*W, F*) of *G* = (*V, E*) is the subgraph for which *x, y* ∈ *W* and (*x, y*) ∈ *E* implies that (*x, y*) ∈ *F*. We call *W* ⊆ *V* a *(strongly) connected component* of *G* = (*V, E*) if *G*[*W*] is an *inclusion-maximal* (strongly) connected subgraph of *G*.

Given a digraph *G* = (*V, E*) and a partition {*V*_1_, *V*_2_, …, *V*_*k*_}, *k* ≥ 1 of its vertex set *V*, the *quotient digraph G/*{*V*_1_, *V*_2_, …, *V*_*k*_} *has as vertex set* {*V*_1_, *V*_2_, …, *V*_*k*_} and two distinct vertices *V*_*i*_ and *V*_*j*_ form an edge (*V*_*i*_, *V*_*j*_) in *G/*{*V*_1_, …, *V*_*k*_} if there are vertices *x* ∈ *V*_*i*_ and *y* ∈ *V*_*j*_ with (*x, y*) ∈ *E*. Note, that edges (*V*_*i*_, *V*_*j*_) in *G/*{*V*_1_, …, *V*_*k*_} do not necessarily imply that (*x, y*) form an edge in *G* for *x* ∈ *V*_*i*_ and *y* ∈ *V*_*j*_. Nevertheless, at least one such edge (*x, y*) with *x* ∈ *V*_*i*_ and *y* ∈ *V*_*j*_ must exist in *G* given that (*V*_*i*_, *V*_*j*_) is an edge in *G/*{*V*_1_, …, *V*_*k*_}

A cycle *C* in a digraph *G* = (*V, E*) of length *n* is an ordered sequence of *n >* 1 (not necessarily distinct) vertices (*v*_1_, …, *v*_*n*_) such that (*v*_*n*_, *v*_1_) ∈ *E* and (*v*_*i*_, *v*_*i*+1_) ∈ *E*, 1 ≤ *i < n*. A digraph that does contain cycles is a *DAG* (directed acyclic graph). Define the relation ⪯_*G*_ of *V* such that *v* ⪯_*G*_ *w* if there is a directed path from *w* to *v*. A vertex *x* is a *parent* of *y* if (*x, y*) ∈ *E*. In this case, *y* is *child* of *x*. Then *G* is DAG if and only if ⪯_*G*_ is a partial order. We write *y* ≺_*G*_ *x* if *y* ⪯_*G*_ *x* and *x*≠ *y*. A *topological order* of *G* is a total order ≪ on *V* such that (*v, w*) ∈ *E* implies that *v* ≪ *w*. It is well known that a digraph *G* admits a topological order if and only if *G* is a DAG. In this case, *x* ≺_*G*_ *y* implies *y* ≪ *x*, i.e., ≪ is a linear extension of ≺. Note that ⪯ and ≪ are arranged in opposite order. The effort to check whether *G* admits a topological order ≪ and, if so, to compute ≪ is linear, i.e., in *O*(|*V* | + |*E*|) [13]. If *C*_1_, …, *C*_*k*_, *k* ≥ 1 are the strongly connected components of a digraph, then *G/*{*C*_1_, *C*_2_, …, *C*_*k*_} is a DAG.

A DAG *G* is *rooted* if it contains a unique ⪯_*G*_-maximal element *ρ*_*G*_ called the *root*. Note that *ρ*_*G*_ is ≪-minimal. A rooted tree *T* with vertex set *V* (*T*) is a DAG such that every vertex *x* ∈ *V* (*T*)\{*ρ*_*T*_ } has a unique parent. The rooted trees considered here do not contain vertices *v* with indeg(*v*) = outdeg(*v*) = 1. A vertex *x* is an *ancestor* of *y* if *y* ⪯_*T*_ *x*, i.e., if *x* is located on the unique path from *ρ*_*T*_ to *y*. A vertex in *T* without a child is a *leaf*. The set of leaves of *T* will be denoted by *L*(*T*). The elements in *V* ^0^(*T*) := *V* (*T*) \ *L*(*T*) are called the inner vertices. We write *T* (*u*) for the subtree of *T* induced by {*v* ∈ *V* (*T*) | *v* ⪯_*T*_ *u*}. Note that *u* is the root of *T* (*u*).

For a subset *W* ⊆ *L*(*T*), the *least common ancestor* lca_*T*_ (*W*) of *W* is the unique ⪯_*T*_ -minimal vertex that is an ancestor of each *w* ∈ *W*. If *W* = {*x, y*}, we write lca_*T*_ (*x, y*) := lca_*T*_ ({*x, y*}). A rooted tree *T* is *ordered*, if the children of every vertex in *T* are ordered. Rooted trees *T*_1_, …, *T*_*k*_, *k* ≥ 2 *are joined under a new root in the tree T* if *T* is obtained by the following procedure: add a new root *ρ*_*T*_ and all trees *T*_1_, …, *T*_*k*_ to *T* and connect the root 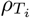 of each tree *T*_*i*_ to *ρ*_*T*_ with an edge (*ρ*_*T*_,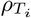).

### Directed Cographs

Di-cographs generalize the notion of undirected cographs [2–4, 6] and are defined recursively as follows: (i) the single vertex graph *K*_1_ is a di-cograph, and (ii) if *G* and *H* are di-cographs, then 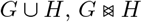, and *G ⊳ H* are di-cographs [8, 16].

Every di-cograph *G* = (*V, E*) *is explained by* an ordered rooted tree *T* = (*W, F*), called a *cotree* of *G*, with leaf set *L*(*T*) = *V* and a (vertex-)labeling function 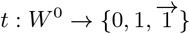 that uniquely determines the set of edges 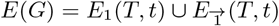 and the set of non-adjacent pairs of vertices *E*_0_(*T, t*) of *G* as follows:

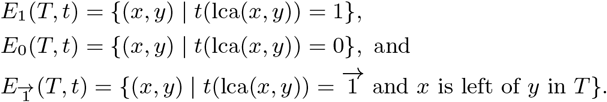

Note that *E*_*i*_(*T, t*) = *E*_*i*_(*T, t*)^sym^ for *i* ∈ {0, 1} since lca(*x, y*) = lca(*y, x*). Every di-cograph *G* = (*V, E*) is explained by a unique *discriminating* cotree (*T, t*) satisfying *t*(*x*)≠ *t*(*y*) for all (*x, y*) ∈ *E*(*T*). Every cotree (*T* ^*′*^, *t*^*′*^) that explains *G* is a “refinement” of its discriminating cotree (*T, t*), i.e., (*T, t*) is obtained by contracting all edges (*x, y*) ∈ *E*(*T*^*′*^) with *t*^*′*^(*x*) = *t*^*′*^(*y*) [1]. Determining whether a digraph is a di-cograph, and if so, computing its discriminating cotree requires *O*(|*V* | + |*E*|) time [2, 8, 15].

## 3 Fitch graphs and Fitch-satisfiability

### 3.1 Basic properties of Fitch graphs

Fitch Graphs are defined in terms of edge-labeled rooted trees *T* with an *edge-labeling λ* : *E* → {0, 1} and leaf set *L*(*T*) = *V*. The graph 𝔾(*T, λ*) = (*V, E*) contains an edge (*x, y*) for *x, y* ∈ *V* if and only if the (unique) path from lca_*T*_ (*x, y*) to *y* contains at least one edge *e* ∈ *E*(*T*) with label *λ*(*e*) = 1. The edge set of 𝔾(*T, λ*) by construction is a binary irreflexive relation on *V*. A directed graph *G* is a *Fitch graph* if there is a tree (*T, λ*) such that *G* ≃ 𝔾(*T, λ*). Fitch graphs form a proper subset of directed cographs [7]. Therefore, they can be explained alternatively by a cotree.

#### Definition 3.1.

*A cotree* (*T, t*) *is a* Fitch-cotree *if there are no two vertices v, w* ∈ *V* ^0^(*T*) *with w* ≺_*T*_ *v such that either (i) t*(*v*) = 0 ≠ *t*(*w*) or *(ii)*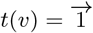 *t*(*w*)=1 and *w* ∈ *V* (*T* (*u*)) *where u is a child of v distinct from the right-most child of v*.

In other words, a Fitch-cotree satisfies:

a. No vertex with label 0 has a descendant with label 1 or 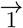.
b. If a vertex *v* has label 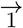, then the subtree *T* (*u*) rooted at a child *u* of *v* – except possibly the right-most one – does not contain vertices with label 1. In particular, if *T* is discriminating, then *T* (*u*) is either a star-tree whose root *u* has label *t*(*u*) = 0 or *u* is a leaf. In either case, the di-cograph *G*[*L*(*T* (*u*))] defined by the subtree *T* (*u*_1_) of left-most child *u*_1_ of *v*, is edge-less.

Fitch graphs have several characterizations that will be relevant throughout this contribution. We summarize [10, L. 2.1] and [7, Theorem 2] in the following statement.

#### Theorem 3.2.

*For every digraph G* = (*V, E*), *the following statements are equivalent*.

1. *G is a Fitch graph*.
2. *G does not contain an induced F*_1_, *F*_2_, …, *F*_8_ *(cf. Figure 1)*.
3. *G is a di-cograph that does not contain an induced F*_1_, *F*_5_ *and F*_8_ *(cf. Figure 1)*.
4. *G is a di-cograph that is explained by a Fitch-cotree*.
5. *Every induced subgraph of G is a Fitch graph, i*.*e*., *the property of being a Fitch graph is hereditary*.

*Fitch graphs can be recognized in O*(|*V* | + |*E*|) *time. In the affirmative case, the (unique least-resolved) edge-labeled tree* (*T, λ*) *can be computed in O*(|*V* |) *time*.

Alternative characterizations can be found in [11]. The procedure `cotree2fitchtree` described in [7] can be used to transform a Fitch cotree (*T, t*) that explains a Fitch graph *G* into an edge-labeled tree (*T* ^*′*^, *λ*) that explains *G* in *O*(|*V* (*T*)|) time, avoiding the construction of the di-cograph altogether.

For later reference, we provide the following simple results.

#### Lemma 3.3.

*The graph obtained from a Fitch graph by removing all bi-directional edges is a DAG*.

*Proof*. Let *G* = (*V, E*) be the graph obtained from the Fitch graph *F* by removing all bi-directional edges and assume, for contradiction, that *G* is not acyclic. Let *C* = (*v*_1_, …, *v*_*n*_) be a shortest cycle in *G* with *n* ≥ 3 vertices. If *n* = 3, then *G* contains *F*_2_ as an induced subgraph, and thus, *F*_2_ is an induced subgraph of *F*. This contradicts Theorem 3.2, as *F* is a Fitch graph. Hence, *n >* 3 must hold. Now consider *v*_1_, *v*_2_ and *v*_3_ on *C* with edges (*v*_1_, *v*_2_), (*v*_2_, *v*_3_) ∈ *E*. Since *G* does not contain cycles on three vertices, we have (*v*_3_, *v*_1_) ∉ *E* and either (*v*_1_, *v*_3_) ∈ *E* or (*v*_1_, *v*_3_) ∉ *E*. If (*v*_1_, *v*_3_) ∈ *E*, then *G* contains a cycle (*v*_1_, *v*_3_, …, *v*_*n*_) shorter than *C*; a contradiction to the choice of *C* as a shortest cycle. Hence, (*v*_1_, *v*_3_) ∉ *E* must hold. This leaves two cases for *F* :

1. (*v*_1_, *v*_3_), (*v*_3_, *v*_1_) ∈ *E*(*F*). In this case *v*_1_, *v*_2_ and *v*_3_ induce an *F*_4_ in *F* ;
2. (*v*_1_, *v*_3_), (*v*_3_, *v*_1_) ∉ *E*(*F*), i.e., *v*_1_ and *v*_3_ are not adjacent in *F*. In this case *v*_1_, *v*_2_ and *v*_3_ induce an *F*_3_ in *F* .

Both alternatives contradict the assumption that *F* is a Fitch graph. Thus, *G* must be acyclic. □

#### Corollary 3.4.

*Every Fitch graph without bi-directional edges is a DAG*.

Removal of the bi-directional edges from the Fitch graph *A*_6_ yields the graph *F*_1_, i.e., although removal of all bi-directional edges from Fitch graphs yields a DAG it does not necessarily result in a Fitch graph.

#### Corollary 3.5.

*Let G be a directed graph without non-adjacent pairs of vertices and without bi-directional edges. Then, G is a Fitch graph if and only if it is a DAG*.

*Proof*. Suppose that *G* is a directed graph without non-adjacent pairs of vertices and without bi-directional edges. Hence, *G* cannot contain the forbidden subgraphs *F*_1_, *F*_3_, …, *F*_8_. If *G* is a DAG, it also cannot contain *F*_2_. Thus, Theorem 3.2 implies that *G* a Fitch graph. Conversely, if *G* is a Fitch graph without bidirectional edges, Corollary 3.4 implies that *G* is a DAG.

#### Lemma 3.6.

*For every Fitch graph F there is a Fitch graph F*^*′*^ *with V* (*F*) = *V* (*F*^*′*^) *and E*(*F*) ⊆ *E*(*F*^*′*^) *that contains no non-adjacent vertices and E*(*F*^*′*^) \ *E*(*F*) *contains no bi-directional edges*.

*Proof*. Let *F* be a Fitch graph. By Lemma 3.3, the graph *G* obtained from *F* by removing all bi-directional edges is a DAG and thus admits a topological order ≪ on *V* (*F*). Construct *F*^*′*^ by adding to *F* the edge (*x, y*) whenever *x* ≪ *y*, (*x, y*) ∉ *E*(*F*) and (*y, x*) ∉ *E*(*F*) for *x, y* ∈ *V* (*F*). The resulting graph *F*^*′*^ contains no non-adjacent vertices and *E*(*F*^*′*^) \ *E*(*F*) contains no bi-directional edges.

It remains to show that *F*^*′*^ is a Fitch graph. Let *x, y, z* be three vertices of *F*. By construction, if *F* [{*x, y, z*}] is isomorphic to *A*_1_, *A*_4_, *A*_5_ or *A*_6_, then *F*^*′*^[{*x, y, z*}] = *F* [{*x, y, z*}]. If *F* [{*x, y, z*}] is isomorphic to one of *A*_2_, *A*_3_ or *A*_8_, then *F*^*′*^[{*x, y, z*}] ≃ *A*_5_ since *F*^*′*^[{*x, y, z*}] is acyclic. Finally, if *F* [{*x, y, z*}] ≃ *A*_7_, then *F*^*′*^[{*x, y, z*}] ≃ *A*_6_. In summary, *F*^*′*^[{*x, y, z*}] does not contain one of the forbidden induced subgraph and thus Theorem 3.2 implies that *F*^*′*^ is a Fitch graph. □

### 3.2 Characterizing Fitch-satisfiability

Throughout we consider 3-tuples of (partial) relations 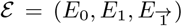 on *V* such that *E*_0_ and *E*_1_ are symmetric and 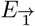 is antisymmetric.

#### Definition 3.7

(*Fitch-sat*). ε *is* Fitch*-satisfiable (in short* Fitch-sat*), if there is a full tuple* 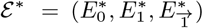 *(with* 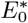 *and* 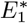 *being symmetric and* 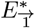 *being antisymmetric) that is explained by a Fitch-cotree* (*T, t*) *and extends* ε .

If ε is *Fitch-sat* with ε^*∗*^ explained by (*T, t*), we say, by slight abuse of notation, that ε is explained by (*T, t*). The problem of finding a tuple ε^*∗*^ that extends ε and that is explained by an arbitrary cotree was investigated in [16]. Theorem 3.2 together with the definition of cotrees and Def. 3.7 implies the following result.

#### Corollary 3.8.

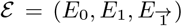 *on V is* Fitch-sat *precisely if it can be extended to a full tuple* 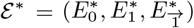 *for which* 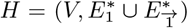 *is a Fitch graph. In particular, there is a discriminating Fitch-cotree that explains* ε, ε^*∗*^, *and H*.

As shown next, Fitch-satisfiability is a hereditary graph property.

#### Lemma 3.9.

*A partial tuple* ε *on V is* Fitch-sat *if and only if* ε [*W*] *is* Fitch-sat *for all W* ⊆ *V* .

*Proof*. The *if* direction follows from the fact that ε [*W*] is *Fitch-sat* for *W* = *V* because ε [*V*] = ε. For the *only-if* direction let *W* ⊆ *V* and assume that 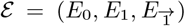 is *Fitch-sat*. By Corollary 3.8, ε can be extended to a full 3-tuple 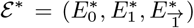 for which 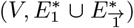 is a Fitch graph. Since the property of being a Fitch graph is a hereditary (Theorem 3.2), 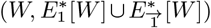 is also a Fitch graph, as 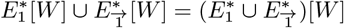. Moreover, we have 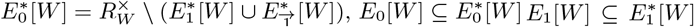,and 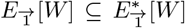, and thus 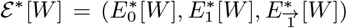 is a full tuple that extends ε [*W*]. In summary, ε [*W*] is *Fitch-sat* . □

For the characterization of *Fitch-sat* tuples in Theorem 3.11, we need the following result.

#### Lemma 3.10.

*Let* 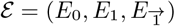 *be a* Fitch-sat *partial tuple on V that is explained by the discriminating Fitch-cotree* (*T, t*) *and let* 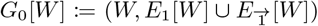 *for W* ⊆ *V*. *If there is a vertex u* ∈ *V* ^0^(*T*) *such that t*(*u*) = 0, *then G*_0_[*C*] *is edge-less for all C* ⊆ *L*(*T* (*u*)).

*Proof*. Let 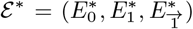 be the full tuple that is explained by the discriminating Fitch-cotree (*T, t*) and that extends the partial tuple 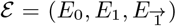 on *V*. Assume that there is a vertex *u* ∈ *V* ^0^(*T*) such that *t*(*u*) = 0. Since (*T, t*) is a Fitch-cotree, there is no inner vertex *v* such that *v* ≺_*T*_ *u* and *t*(*v*)≠ 0. Let *W* = *L*(*T* (*u*)). Therefore, all vertices *v* ≺_*T*_ *u* must be leaves, as (*T, t*) is discriminating. This implies that for all distinct *x, y* ∈ *W* it holds that lca(*x, y*) = *u* and, thus, *t*(lca(*x, y*)) = *t*(*u*) = 0. Since (*T, t*) explains ε^*∗*^ it follows that 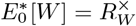 and, thus, 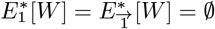. Since 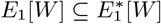 and 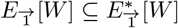, we have 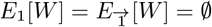. Hence, 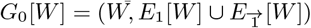 is edge-less. Trivially, *G*_0_[*C*] must be edge-less for all *C* ⊆ *W* . □

We are now in the position to provide a characterization of *Fitch-sat* partial tuples.

#### Theorem 3.11.

*The partial tuple* 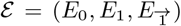 *on V is* Fitch-sat *if and only if at least one of the following statements holds*.

(S1) 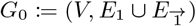*is edge-less*.
(S2) (a) 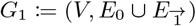is disconnected and (b) *ε* [*C*] is *Fitch-sat* for all connected components *C* of *G*_1_
(S3) (a)
  I. 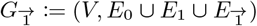 contains *k >* 1 strongly connected components *C*_1_, …, *C*_*k*_ collected in 𝒞 and
  II. there is a *C* ∈ 𝒞 for which the following conditions are satisfied:
    i. *G*_0_[*C*] is edge-less.
    ii. *C* is ≪-minimal for some topological order ≪ on 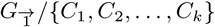
ε [*V* \ *C*] is *Fitch-sat* .

*Proof*. We start with proving the *if* direction and thus assume that 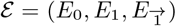 is a partial tuple on *V* that satisfies at least one of (S1), (S2) and (S3).

First assume that *ε* satisfies (S1). Hence, 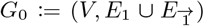 is edge-less and we can replace *ε* _0_ by 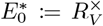 to obtain the full tuple 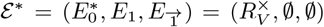. One easily verifies that the star tree with leaf-set *V* and whose root is labeled “0” explains *ε* ^*∗*^ and is, in particular, a Fitch-cotree. Hence, ε ^*∗*^ is *Fitch-sat* .

Next assume that ε satisfies (S2). Hence, 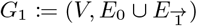 is disconnected and *ε*[*C*_*i*_] is *Fitch-sat* for each connected component *C*_1_, …, *C*_*k*_, *k >* 1. Thus, we can extend each *ε* [*C*_*i*_] to a full sat *ε* ^*∗*^[*C*_*i*_] that is explained by a Fitch-cotree (*T*_*i*_, *t*_*i*_). Let 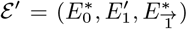 be the resulting partial tuple obtained from *ε* such that 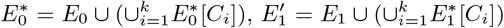, and 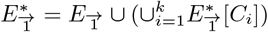. To obtain 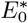 and 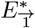, we added only pairs (*x, y*) with *x, y* ∈ *C*_*i*_ to *E*_0_ and 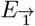, respectively. Therefore, 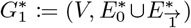 remains disconnected and has, in particular, connected component *C*_1_, …, *C*_*k*_. We now extend ε ^*′*^ to a full tuple ε ^*∗*^ by adding all 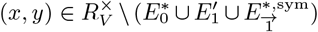 to 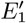 to obtain 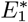. Note that the pairs in 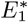 do not alter disconnectedness of 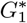, i.e., 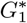 has still connected components *C*_1_, … *C*_*k*_. By construction, we have 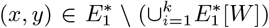 with 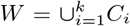 if and only if *x* and *y* are in distinct connected components of 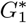. We now join the Fitch-cotrees (*T*_1_, *t*_1_), … (*T*_*k*_, *t*_*k*_) under a common root *ρ* with label “1” and keep the labels of all other vertices to obtain the cotree (*T, t*). Since each (*T*_*i*_, *t*_*i*_) is a Fitch-cotree, and since no constraints are imposed on vertices with label “1” (and thus on the root *ρ* of *T*), it follows that (*T, t*) is a Fitch-cotree. Moreover, by construction, for all *x* and *y* that are in distinct connected components of *G*^*∗*^ we have lca_*T*_ (*x, y*) = *ρ* and thus *t*(lca_*T*_ (*x, y*)) = 1. Hence, 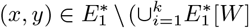 if and only of lca_*T*_ (*x, y*) = *ρ* and thus *t*(lca_*T*_ (*x, y*)) = 1. Moreover, each Fitch-cotree (*T*_*i*_, *t*_*i*_) explains ε ^*∗*^[*C*_*i*_]. Taking the latter arguments together, the Fitch-cotree (*T, t*) explains the full tuple ε ^*∗*^, and thus ε is *Fitch-sat* .

Finally, assume that ε satisfies (S3). By (S3.a.I), 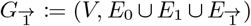 has *k >* 1 strongly connected components *C*_1_, …, *C*_*k*_ collected in 𝒞. Moreover, by (S3.a.II.i), there is a *C* ∈ 𝒞 for which *G*_0_[*C*] is edge-less, where *C* is ≪-minimal for some topological order ≪ on 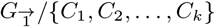 and where *ε* [*Ĉ*] is *Fitch-sat* for *Ĉ* = *V* \ *C*. Since *ε* [*Ĉ*]is *Fitch-sat, ε* [*Ĉ*] can be extended to a full tuple 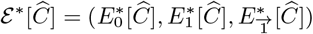 that is a explained by a Fitch-cotree 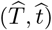. Moreover, *G*_0_[*C*] is edge-less and thus 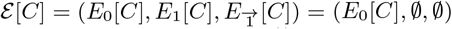. We can now extend ε [*C*] to a full tuple 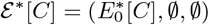 by adding all pairs 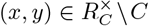 to *E*_0_[*C*]. Clearly, 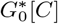 remains edge-less. Hence, ε ^*∗*^[*C*] is explains by the Fitch-cotree (*T* ^*′*^, *t*^*′*^) where *T* ^*′*^ is a star-tree whose root 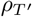 has label (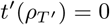) We finally join (*T* ^*′*^, *t*^*′*^) and 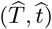 under a common root *ρ*_*T*_ to obtain the cotree (*T, t*) where we put 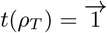 and assume that 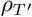 is placed left of 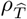. The latter, in particular, ensures that (*T, t*) is a Fitch-cotree.

It remains to show that the full-set ε ^*∗*^ explained by (*T, t*) extends ε. First, observe that, by construction, 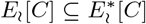 and 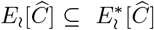 for 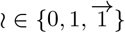. By construction of (*T, t*), we have lca (*x, y*) = *ρ*_T_ for all *x* ∈ *C* and 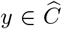. By construction, (*T, t*) explains ε ^*∗*^[*C*] and ε ^*∗*^[*Ĉ*] and thus, ε ^*∗*^[*C*] extends ε [*C*] while and ε ^*∗*^[*Ĉ*] extends ε [*Ĉ*].

Hence, it remains to consider all vertices *x* ∈ *C* and *y* ∈ *Ĉ*. Let *x* ∈ *C* and *y* ∈ *Ĉ*. Since *C* is ≪-minimal for some topological order ≪ on 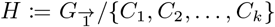, there is no edge (*C*^*′*^, *C*) in *H* for any *C*^*′*^ ∈ 𝒞 with *C*^*′*^ ⊆ *Ĉ*. Hence, there is no edge (*y, x*) in 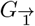. Therefore, any edge between *x* and *y* that is contained in a relation of ε must satisfy 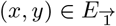 (since *E*_0_ and *E*_1_ are symmetric). Since 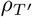 is placed left of 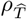, it follows that *x* ∈ *C* is placed left of *y* ∈ *C*^*′*^ for every *C*^*′*^ ⊆ *Ĉ*. This, together with the fact that lca_*T*_ (*x, y*) = *ρ*_*T*_ and 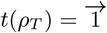, implies that 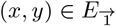 is correctly explained for all *x* ∈ *C* and *y* ∈ *C*. Moreover, for every *x* ∈ *C* and *y* ∈ *Ĉ*, we have by construction of (*T, t*) that 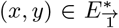 and 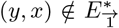. Hence, 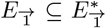 holds. Thus, the full-set ε ^*∗*^ that is explained by (*T, t*) extends ε and, therefore, ε is *Fitch-sat*.

For the *only-if* direction, we assume that the partial tuple 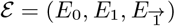 on *V* is *Fitch-sat*. If |*V* | = 1, then (S1) is obviously satisfied. We therefore assume |*V* | ≥ 2 from here on. Let 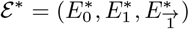 be a full tuple that is *Fitch-sat* and that extends ε. By Corollary 3.8, there is a discriminating Fitch-cotree (*T, t*) endowed with the sibling order order *<* that explains ε and ε ^*∗*^ and has leaf set *V*. Moreover, the root *ρ* of *T* has at *r* ≥ 2 children *v*_1_, …, *v*_*r*_ ordered from left to right according to *<* and has one of the three labels 0, 1, 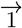. In the following, *L*_*i*_ = {*x* ∈ *L*(*T*) | *x* ⪯ *v*_*i*_} denotes the set of all leaves *x* of *T* with *x* ⪯ *v*_*i*_. We make frequent use of the fact that lca(*x, y*) = *ρ* and thus, *t*(lca(*x, y*)) = *t*(*ρ*) for all *x* ∈ *L*_*i*_, *y* ∈ *L*_*j*_ with *i*≠ *j* without further notice. We proceed by considering the three possible choices of *t*(*ρ*).

If *t*(*ρ*) = 0, then Lemma 3.10 implies that 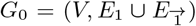 is edge-less and thus ε satisfies (S1). If *t*(*ρ*) = 1, then *t*(*ρ*) = *t*(lca(*x, y*)) = 1 for all *x* ∈ *L*_*i*_, *y*∈ *L*_*j*_ with *i* ≠ *j*. Hence, 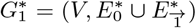 must be disconnected. Since 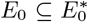 and 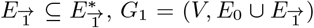 is a subgraph of 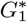 and thus, *G*_1_ is also disconnected. For each of the connected components *C* of *G*_1_, Lemma 3.9 implies that ε [*C*] is *Fitch-sat*. Therefore, ε satisfies (S2).

If 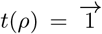 we have 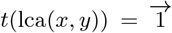 for all *x* ∈ *L*_*i*_, *y* ∈ *L*_*j*_ with *i*≠ *j*. Since (*T, t*) explains ε, we observe that 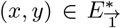 and 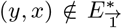 for all *x* ∈ *L*_*i*_, *y* ∈ *L*_*j*_ with 1 ≤ *i < j* ≤ *r*. Thus, 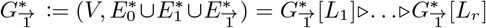. Hence, 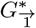 contains more than one strongly connected component (which may consist of a single vertex). In particular, each strongly connected component of 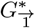 must be entirely contained in some *L*_*i*_, *i* ∈ {1, …, *r*}. Let 𝒞 be the set of strongly connected components of 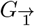. Since 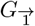 is a subgraph of 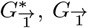, contains more than one strongly connected component and therefore, ε satisfies (S3.a.I). In particular, each *C* ∈ 𝒞 must be contained in some strongly connected component of 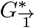. Taken the latter arguments together, each *C* ∈ 𝒞 must satisfy *C* ⊆ *L*_*i*_ for some *i* ∈ {1, …, *r*}.

Consider now the left-most child *v*_1_ of *ρ*. Since (*T, t*) is a discriminating Fitch-cotree, this vertex *v*_1_ is either a leaf of *T* or *t*(*v*_1_) = 0. In either case, *G*_0_[*L*_1_] is edge-less (cf. Lemma 3.10). By the latter arguments and since *L*_1_≠ ∅, there is some *C* ∈ 𝒞 with *C* ⊆ *L*_1_. Since *G*_0_[*C*] ⊆ *G*_0_[*L*_1_] is edge-less, Condition (S3.a.II.i) holds.

Let *C*^*′*^ ∈ 𝒞 \ {*C*}. Suppose first that *C*^*′*^ ⊆ *L*_1_. Assume, for contradiction, that there are edges (*x, y*) or (*y, x*) with *x* ∈ *C* and *y* ∈ *C*^*′*^ in 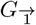. Since *G*_0_[*L*_1_] is edge-less, we have (*x, y*) ∈ *E*_0_ or (*y, x*) ∈ *E*_0_. Since *E*_0_ is symmetric (*x, y*) ∈ *E*_0_ implies (*y, x*) ∈ *E*_0_ and *vice versa*. Hence, the union *C* ∪ *C*^*′*^ would be strongly connected in 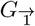 ; a contradiction since *C* and *C*^*′*^ are distinct. Hence, there cannot be any edge between vertices in *C* and *C*^*′*^ in 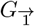 whenever *C*^*′*^ ⊆ *L*_1_. Suppose now that *C*^*′*^ ⊆ *L*_*i*_, 1 *< i* ≤ *r*. As argued above, 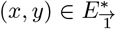 and 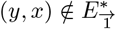 for all *x* ∈ *C* ⊆ *L*_1_ and *y* ∈ *C*^*′*^ ⊆ *L*_*i*_. Since 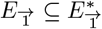, any to adjacent vertices *x* ∈ *C* and *y* ∈ *C*^*′*^ in 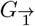 must satisfy 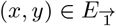 and 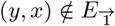.

Let 𝒞 = {*C*_1_, …, *C*_*k*_}, *k >* 1. Hence, the quotient 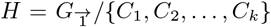 is a DAG. By the latter arguments, there are no edges (*y, x*) with *x* ∈ *C* and *y* ∈ *C*^*′*^ for any *C*^*′*^ ∈ 𝒞 \ {*C*}. Hence, there is no edge (*C*^*′*^, *C*) in *H* for any *C*^*′*^ ∈ 𝒞 \ {*C*}. Since *H* is a DAG, it admits a topological order ≪ such that *C* is ≪-minimal. Hence, (S3.a.II.ii) is satisfied. Finally, Lemma 3.9 implies that ε [*V* \ *C*] is *Fitch-sat* and thus (S3.b) is satisfied. In summary, ε satisfied (S3).

## 4 Recognition Algorithm and Computational Complexity

The proof of Theorem 3.11 provides a recipe to construct a Fitch-cotree (*T, t*) explaining a tuple ε. We observe, furthermore, that two or even all three alternatives (S1), (S2.a), and (S3.a) may be satisfied simultaneously, see Figure 2 for an illustrative example. In this case, it becomes necessary to check stepwisely whether conditions (S2.b) and/or (S3.b) holds. Potentially, this result in exponential effort to determine recursively whether ε [*C*] or ε [*V* \ *C*] is *Fitch-sat*. The following simple lemma shows, however, that the alternatives always yield consistent results:

**Figure 2:**
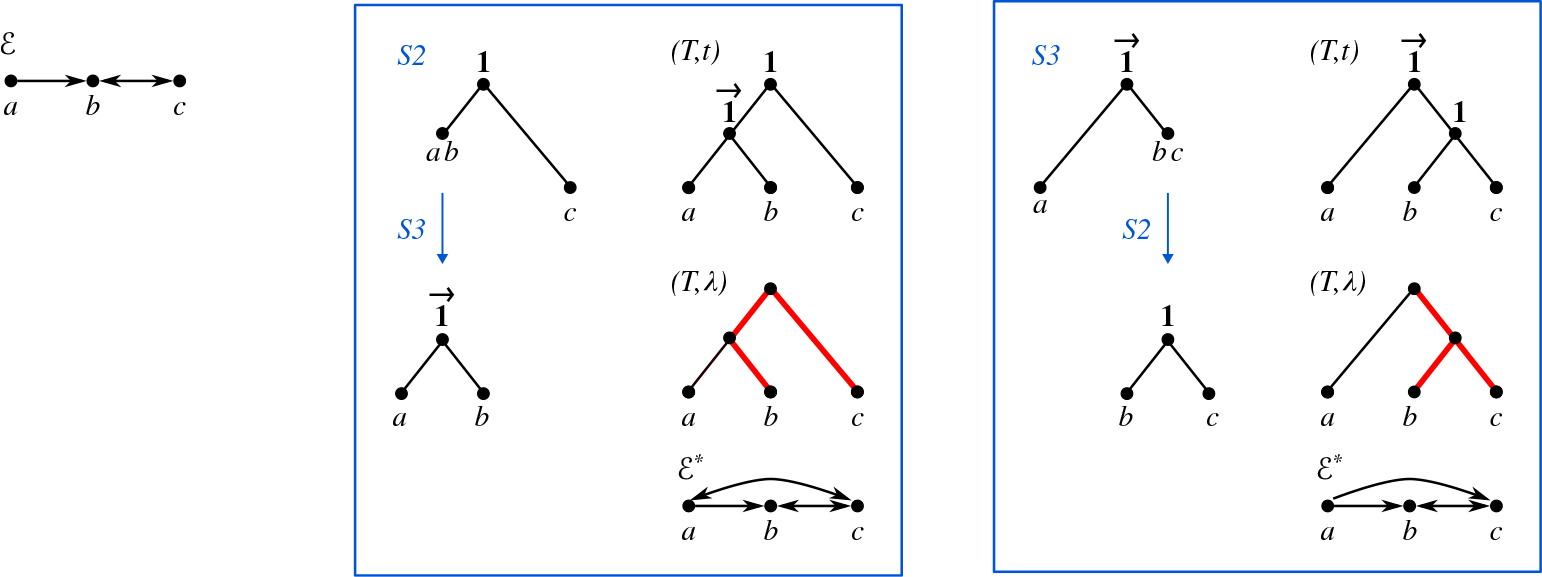
A partial tuple 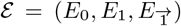 on *V* = {*a, b, c*} with *E*_0_ = ∅, *E*_1_ = {(*b, c*), (*c, b*)} (bidirectional arc), and 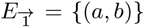 .(single arc) is shown on the left. Observe that (S1) is not satisfied while (S2.a) and (S3.a) hold for ε. Application of the different rules and subsequent construction of the Fitch-cotrees that explain the subgraphs induced by the respective (strongly) connected components results in two Fitch-cotrees that both explain ε. Hence, we obtain two different edge-labeled Fitch-trees (*T, λ*) (with HGT-edges drawn in bold-red) that both explain ε .

### Lemma 4.1.

*Let* ε = (*E*_0_, *E*_1_, 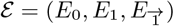) *be a partial tuple on V*. *Then*

> (S1) and (S2.a) implies (S2.b);
>
> (S1) and (S3.a) implies (S3.b);
>
> (S2a) and (S3a) implies that (S2.b) and (S3.b) are equivalent.

*Proof*. If (S1) holds, then Theorem 3.11 implies that ε is *Fitch-sat*. If (S2.a) holds, then heredity (Lemma 3.9) implies that ε [*C*] is *Fitch-sat* and thus (S2.b) is satisfied. Analogously, if (S3.a) holds, then ε [*V* \ *C*] is *Fitch-sat* and thus (S3.b) holds. Now suppose (S2a) and (S3a) are satisfied, but (S1) does not hold. Then ε is *Fitch-sat* if and only if one of (S2.b) or (S3.b) holds; in the affirmative case, heredity again implies that both (S2.b) and (S3.b) are satisfied. □

### Algorithm 1

Recognition of *Fitch-sat* partial tuple *E* on *V* and reconstruction of a cotree (*T, t*) that explains *E* .

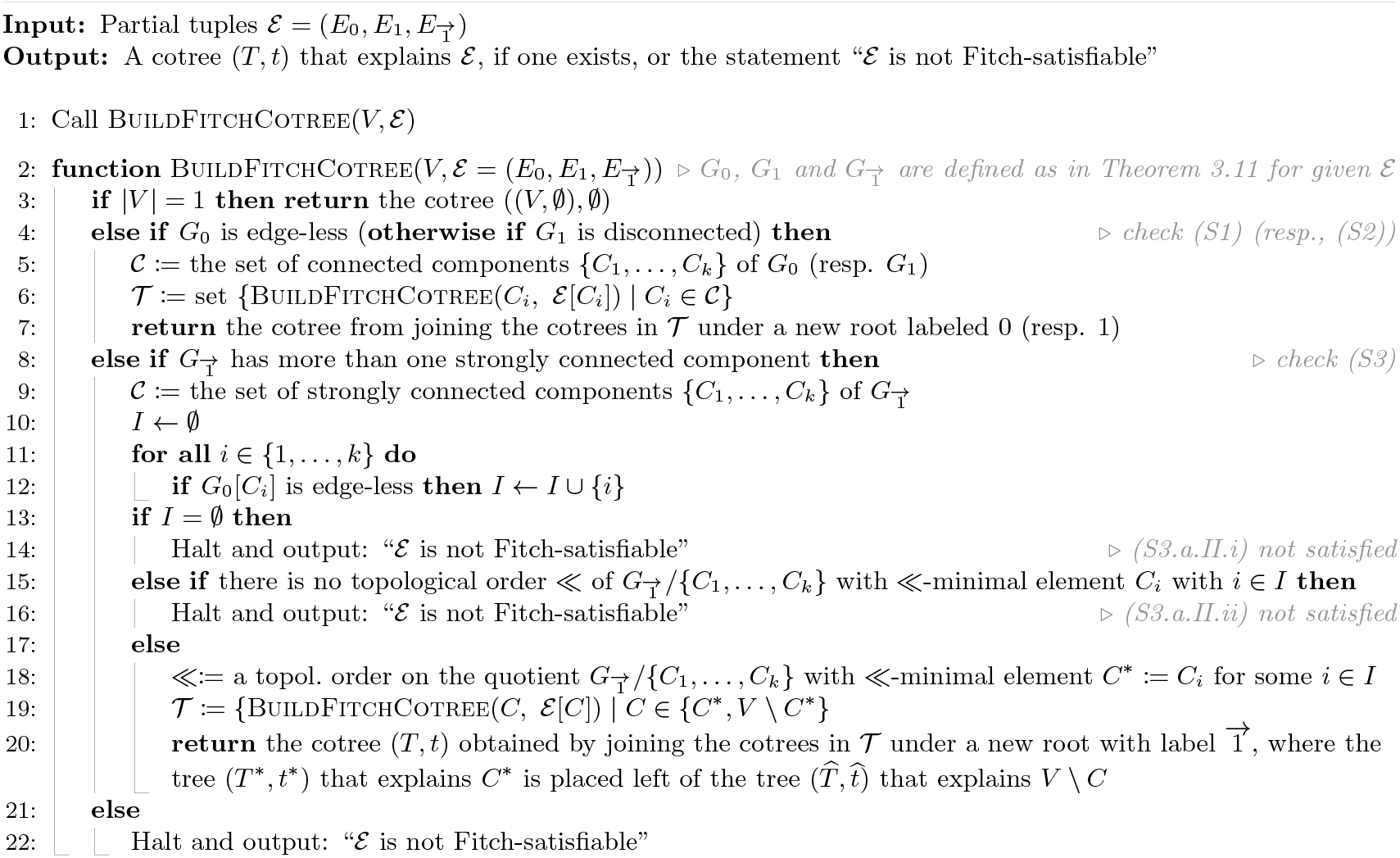

It follows that testing whether ε can be achieved by checking if any one of the three conditions (S1), (S2), or (S3) holds and, if necessary, recursing down on ε [*C*] or ε [*V* \ *C*]. This gives rise to Algorithm 1.

### Lemma 4.2.

*Let* 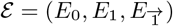 *be a partial tuple. Then Algorithm 1 either outputs a Fitch-cotree* (*T, t*) *that explains* ε *or recognizes that* ε *is not* Fitch-sat.

*Proof*. Let 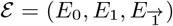 be a partial tuple on *V* that serves as input for Algorithm 1. In Algorithm 1, the function BuildFitchCotree is called, and the algorithm we will try and build a Fitch-cotree explaining a full *Fitch-sat* tuple that extends ε. By Lemma 4.1, the order in which (S1), (S2), and (S3) are tested is arbitrary. If |*V* | = 1, then ε is already full and *Fitch-sat*, and (*T, t*) = ((*V*, ∅); ∅) is a valid Fitch-cotree explaining ε, and thus returned on Line 3. If none of the conditions (S1), (S2), or (S3) is satisfied, Theorem 3.11 implies that ε is not *Fitch-sat*, and Algorithm 1 correctly returns “ε is not *Fitch-sat*”.

If Rule (S1) or (S2.a), resp., is satisfied (Line 4), then Algorithm 1 is called recursively on each of the connected components defined by *G*_0_ or *G*_1_, respectively. In the case of (S1), the connected components are all single vertices, and we obtain a star-tree (*T, t*) whose root is labeled “0” as outlined in the proof of the *if* direction of Theorem 3.11. Since (*T, t*) is a Fitch-cotree that explains ε, it follows that ε is *Fitch-sat*. In the case of (S2.a), condition (S2.b) must be tested. If each of the connected components are indeed *Fitch-sat*, the obtained Fitch-cotrees are joined into a single cotree (*T, t*) that explains a full tuple ε ^*∗*^, which extends ε. In particular, since no constraints are imposed on the root *ρ*_*T*_, which has label “1”, (*T, t*) is a Fitch-cotree. Consequently, ε is *Fitch-sat* and explained by (*T, t*).

Suppose, finally, that ε satisfies (S3.a) and let 𝒞 := {*C*_1_, …, *C*_*k*_} be the the set of strongly connected components of 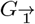. Since (S3.a.II) is satisfied, there must be a *C*_*i*_ ∈ 𝒞 for which *G*[*C*_*i*_] is edge-less, i.e., the set *I* as computed in line 12 cannot be empty. In particular, there is a topological order ≪ of 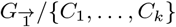 with ≪-minimal element *C*^*∗*^ := *C*_*i*_ for some *i* ∈ *I*. It is now checked, if ε [*C*^*∗*^] and ε [*Ĉ*] are *Fitch-sat*, where *Ĉ* := *V* \ *C*. If this is the case, two Fitch-cotrees (*T*^*∗*^, *t*^*∗*^) and (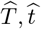) that explain ε [*C*^*∗*^] and ε [*Ĉ*] are returned.

We then join these cotrees in under a new root with label 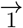, where the tree (*T*^*∗*^, *t*^*∗*^) is placed left from (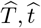). The latter results in a cotree (*T, t*) that explains ε. It remains to show that (*T, t*) is a Fitch-cotree.

Since, *ρ*_*T*_ has precisely two children *ρ*_*T*_*∗* and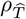, and *ρ*_*T*_*∗* is left of 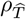, we must show that the tree (*T* ^*∗*^, *t*^*∗*^) computed in the recursive calls does not contain an inner vertex *v* with label *t*(*v*) = 1. The recursive call with ε [*C*^*∗*^] first checks if |*C*^*∗*^| = 1 (line 3) and then continues with checking if *G*_0_[*C*^*∗*^] is edge-less (line 4). Since *G*_0_[*C*^*∗*^] is edge-less, either the single-vertex tree or a star tree whose root has label “0” will be returned. Consequently, (*T*^*∗*^, *t*^*∗*^) does not contain an inner vertex *v* with label *t*(*v*) = 1 and thus, (*T, t*) is a Fitch-cotree that explains ε. Hence, ε is *Fitch-sat* . □

### Theorem 4.3.

*Let* 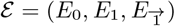 *be a partial tuple, n* = |*V* | *and* 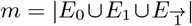. *Then, Algorithm 1 computes a Fitch-cotree* (*T, t*) *that explains* ε *or identifies that* ε *is not* Fitch-sat *in O*(*n*^2^ + *nm*) *time*.

*Proof*. Correctness is established in Lemma 4.2. We first note that in each single call of BuildFitchCotree, all necessary di-graphs defined in (S1), (S2) and (S3) can be computed in *O*(*n* + *m*) time. Furthermore, each of the following tasks can be performed in *O*(*n* + *m*) time: finding the (strongly) connected components of each digraph, construction of the quotient graphs, and finding the topological order on the quotient graph using Kahn’s algorithm [13]. Moreover, the vertex end edge sets *V* [*C*] and ε [*C*] for the (strongly) connected components *C* (or their unions) can be constructed in *O*(*n*+*m*) time by going through every element in *V* and ε and assigning each pair to their respective induced subset. Thus, every pass of BuildFitchCotree takes *O*(*n* + *m*) time. Since every call of BuildFitchCotree either halts or adds a vertex to the final constructed Fitch-cotree, and the number of vertices in this tree is bounded by the number *n* of leaves, it follows that BuildFitchCotree is called at most *O*(*n*) times resulting in an overall running time *O*(*n*(*n* + *m*)). □

## 5. Optimization and Computational Complexity

Instead of asking only for the existence of a Fitch-completion of a tuple 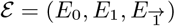, it is of interest to ask for the completion that maximizes a total score for the pairs of distinct vertices *x, y* that are not already classified by ε, i.e.,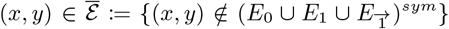. For every pair of vertices *x* and *y* there are four possibilities *x* :: *y* ∈ {*x* ⇌ *y, x* → *y, x* ← *y, x y*} depending on whether *x* and *y* are connected via bi-directional arcs (⇌), only uni-directional arcs (←, →) or if they are not adjacent at all 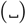. The score *w*(*x* :: *y*) may be a log-odds ratio for observing one of the four possible xenology relationship as determined from experimental data. Let us write 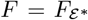 for the Fitch graph defined by the extension ε ^*∗*^ of ε and associate with it the total weight of relations added, i.e.,

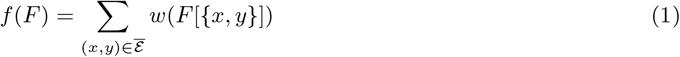

It is then of particular interest to find a Fitch graph 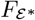 that maximizes *f* over all possible Fitch-completions of ε. The weighted Fitch-completion problem can also be seen as a special case of the problem with the empty tuple ε ^*∅*^ := (∅, ∅, ∅). To see this, suppose first that an arbitrary partial input tuple ε together with weights *w*(*x* :: *y*) for all four possibilities of :: and for all distinct *x, y* with 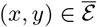 is given. Moreover, we may assume w.l.o.g. that ε can be extended to ε ^*∗*^ such that 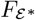 is a Fitch graph (verifying the latter can be done on polynomial-time by Theorem 4.3 and, in the negative case, we can skip any try to extend ε to a Fitch-graph). For each two vertices *x, y* for which 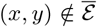 the induced graphs *F* [{*x, y*}] is well-defined, and we extend the weight function to all pairs of vertices by setting, for all 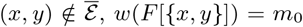 and *w*(*x* :: *y*) = −*m*_0_ for (*x* :: *y*)≠ *F* [{*x, y*}], where 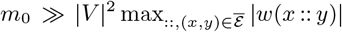. Now consider the weighted Fitch-completion problem with this weight function and an empty tuple ε ^*∅*^. The choice of weights ensures that any Fitch graph *F*^*′*^ maximizing *f* (*F*^*′*^) induces *F*^*′*^[{*x, y*}] = *F* [{*x, y*}] for all pairs {*x, y*} in the input tuple, because not choosing *F* [{*x, y*}] reduces the score by 2*m*_0_ while the the total score of all pairs not specified in the input is smaller than *m*_0_. In order to study the complexity of this task, it therefore suffices to consider the following decision problem.

### Problem 5.1

(Fitch Completion Problem (FC)).

Input: *A set V, an assignment of four weights w*(*x* :: *y*) *to all distinct x, y* ∈ *V where* :: ∈ {⇌, →, ←, }, *and an integer k* ≥ 0.

Question: *Is there a Fitch graph F* = (*V, E*) *such that* 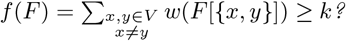

For the NP-hardness reduction, we use the following NP-complete problem [14]

### Problem 5.2

(Maximum Acyclic Subgraph Problem (MAS)).

*G* Input: *A digraph G* = (*V, E*) *and an integer k* ≥ 0.

Question: *Is there a subset E*^*′*^ ⊆ *E such that* |*E*^*′*^| ≥ *k and* (*V, E*^*′*^) *is a DAG?*

### Theorem 5.3.

FC *is NP-complete*.

*Proof*. We claim that FC is in NP. To see this, let *F* = (*V, E*) be a given digraph. We can check whether *f* (*F*) ≥ *k* in polynomial time by iterating over all edges in *F*. In addition, by Theorem 3.2, we can check whether *F* is a Fitch graph in polynomial time by iterating over all 3-subsets of *V* and verifying that none of the induced a forbidden subgraph of Fitch graphs.

To prove NP-hardness, we must show that there is a polynomial-time reduction transforming arbitrary instances *I* of MAS to an instance *I*^*′*^ of FC such that *I* is a yes-instance of MAS if and only if *I*^*′*^ is a yes-instance of FC. Let (*G* = (*V, E*), *k*) be an instance of MAS. The instance (*V, k, w*) of FC uses the following weights for all distinct *x, y* ∈ *V* :

i. If (*x, y*) ∈ *E*, then put *w*(*x* → *y*) = 1
ii. If (*x, y*) ∉ *E*, then put *w*(*x* → *y*) = 0
iii. Put *w*(*x y*) = 0 and *w*(*x* ⇌ *y*) = −|*V* |^2^.

It is easy to verify that the constructed instance of FC can be derived in polynomial time. Note that Condition (i) ensures that, for all *x, y* ∈ *V*, we have *w*(*y* → *x*) = 1 if (*y, x*) ∈ *E* and *w*(*x* → *y*) = *w*(*y* → *x*) = 1 whenever both (*x, y*) and (*y, x*) are edges in *G*. Suppose first that (*G* = (*V, E*), *k*) is a yes-instance of MAS. i.e., there is a subset *E*^*′*^ ⊆ *E* such that |*E*^*′*^| ≥ *k* and *G*^*′*^ = (*V, E*^*′*^) is a DAG. Hence, for any *x, y* ∈ *V* not both (*x, y*) and (*y, x*) can be contained in *E*^*′*^. This, together with the construction of the weights, implies that *f* (*G*^*′*^) = |*E*^*′*^| ≥ *k*. We now extend *G*^*′*^ to a Fitch graph *F*. To this end, observe that *G*^*′*^ admits a topological order ≪. We now add for all pairs *x, y* with *x* ≪ *y* and (*x, y*) ∉ *E*^*′*^ the edge (*x, y*) to obtain the di-graph *F*. Clearly, ≪ remains a topological order of *F* and, therefore, *F* is a DAG. This with the fact that *F* does not contain bi-directional edges or non-adjacent vertices together with Corollary 3.5 implies that *F* is a Fitch graph. In particular, *f* (*F*) ≥ *f* (*G*^*′*^) ≥ *k* and therefore, we obtain a yes-instance of FC.

Assume now that (*V, k, w*) is a yes-instance of FC and thus, there is a Fitch graph *F* such that *f* (*F*) ≥ *k*. Since *w*(*x* ⇌ *y*) = −|*V* |^2^, and the weight for any uni-directed edge and every pair of non-adjacent vertices is 0 or 1 and the maximum number of edges in *F* is 2 · 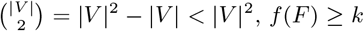. implies that *F* cannot contain bidirectional edges because *k* is non-negative. By Corollary 3.4, *F* is acyclic. Now, take the subgraph *G*^*′*^ of *F* that consists of all edges with weight 1. Clearly, *G*^*′*^ remains acyclic and *f* (*F*) = *f* (*G*^*′*^) ≥ *k*. By construction of the weights, all edges of *G*^*′*^ must have been contained in *G*, and thus, *G*^*′*^ ⊆ *G* is an acyclic subgraph of *G* containing at least *k* edges.

### Problem 5.4

(FC without bi-directional edges).

Input: *A set V, an assignment of four weights w*(*x* :: *y*) *to all distinct x, y* ∈ *V where* :: ∈ {⇌, →, ←, }, *and an integer k*.

Question: *Is there a Fitch graph F* = (*V, E*) *such that* 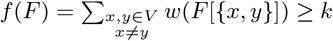 *and, for all x, y* ∈ *V*, (*x, y*) ∈ *E implies* (*y, x*) ∉ *E ?*

Closer inspection of the proof of Theorem 5.3 shows that, for both directions, the respective Fitch graph does not contain bi-directional edges. Hence, we immediately obtain

### Theorem 5.5.

FC without bi-directional edges *is NP-complete*.

Lemma 3.6 ensures the existence of Fitch graphs without non-adjacent vertices. For this family of graphs, we consider

### Problem 5.6

(FC without non-adjacent vertices).

Input: *A set V, an assignment of four weights w*(*x* :: *y*) *to all distinct x, y* ∈ *V where* :: ∈ {⇌, →, ←, }, *and an integer k*.

Question: *Is there a Fitch graph F* = (*V, E*) *such that* 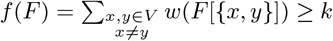 *and, for all x, y* ∈ *V it holds that* (*x, y*) ∈ *E or* (*y, x*) ∈ *E ?*

### Theorem 5.7.

FC without non-adjacent vertices *is NP-complete*.

*Proof*. Containment in NP as well as construction of the instance (*V, k, w*) of FC and the proof of the *only-if* direction is precisely as in the proof of Theorem 5.3. In particular, the proof of the *only-if* direction in Theorem 5.3 shows that the resulting Fitch graph does not contain non-adjacent edges. In addition, if we have a yes-instance (*V, k, w*) of FC and therefore, a Fitch graph *F* with *f* (*F*) ≥ *k* then there is, by Lemma 3.6, in particular a Fitch graph *F*^*′*^ with *f* (*F*) = *f* (*F*^*′*^) whenever *w*(*x y*) = 0. We can now reuse exactly the same arguments as in the *if* direction in Theorem 5.3 to conclude that the subgraph *G*^*′*^ of *F*^*′*^ that consists of all edges with weight 1 is an acyclic subgraph of *G* containing at least *k* edges. Hence, FC without non-adjacent vertices is NP-hard. □

### Problem 5.8

(Fitch Completion Problem with partial Input, FCpI).

Input: *A set V, a partial Fitch-satisfiable tuple* 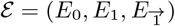 *such that at least one* 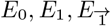. *is non-empty, an assignment of four weights w*(*x* :: *y*) *to all distinct x, y* ∈ *V where* :: ∈ {⇌, →, ←, }, *and an integer k* ≥ 0.

Question: *Is there a Fitch graph F* = (*V, E*) *such that* 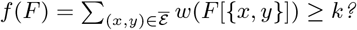

The arguments preceding Problem 5.1 show that FCpI can be seen as a special instance of FC. Hence, these arguments do no imply that FCpI is NP-hard. To address the complexity of FCpI, observe first that that none of the forbidden subgraphs *F*_1_, …, *F*_8_ contain a vertex *w* such that 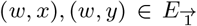. This immediately implies

### Observation 5.9.

*Let G* = (*V, E*) *be a digraph and G*^+*z*^ = (*V* ^*′*^, *E*^*′*^) *be obtained from G by adding a new vertex z to V (resulting in V* ^*′*^*) and the additional arcs* (*z, x*) *to E for all z* ∈ *V (resulting in E*^*′*^*). Then, G is a Fitch-graph if and only if G*^+*z*^ *is a Fitch-graph*.

### Theorem 5.10.

FCpI *is NP-complete*.

*Proof*. FCpI is in NP by similar arguments that show that FC is in NP. Let (*V, k, w*) be an instance of FC. We construct an instance (*V* ^*′*^, ε, *k, w*^*′*^) of FCpI by adding an additional vertex *z* to *V* (resulting in *V* ^*′*^) and the arc (*z, x*) to 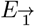 for all *x* ∈ *V* (resulting in 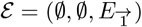)) while keeping the weights *w* for all *x* :: *y* with *x, y* ∈ *V* and adding arbitrary weights to *z* :: *x* for all *x* ∈ *V* and all :: ∈ {⇌, →, ←, }.

Then, (*V, k, w*) is a yes-instance of FC if and only if there is a Fitch graph *F* such that *f* (*F*) ≥ *k* (and thus *f* (*F*) = *f* (*F* ^+*z*^) ≥ *k*) which is, by Observation 5.9 and the choice of the weights, if and only there is a Fitch graph *F*^*′*^ := *F* ^+*z*^ such that *f* (*F*^*′*^) ≥ *k* if and only if (*V* ^*′*^, ε, *k, w*^*′*^) is a yes-instance of FCpI. Hence, FCpI is in NP-hard. □

The structure of the Fitch completion problems suggests a canonical greedy heuristic, in which the weights for the possible 2-vertex graphs are sorted in descending order. For each proposed insertion of *x* :: *y*, we take the one with the hightest weight and for which the extended tupled ε is *Fitch-sat*. The latter can be tested with Algorithm 1. The polynomial-time greedy heuristic is provided in Algorithm 1.

### Algorithm 2

Greedy to compute full *E* that is *Fitch-sat* .

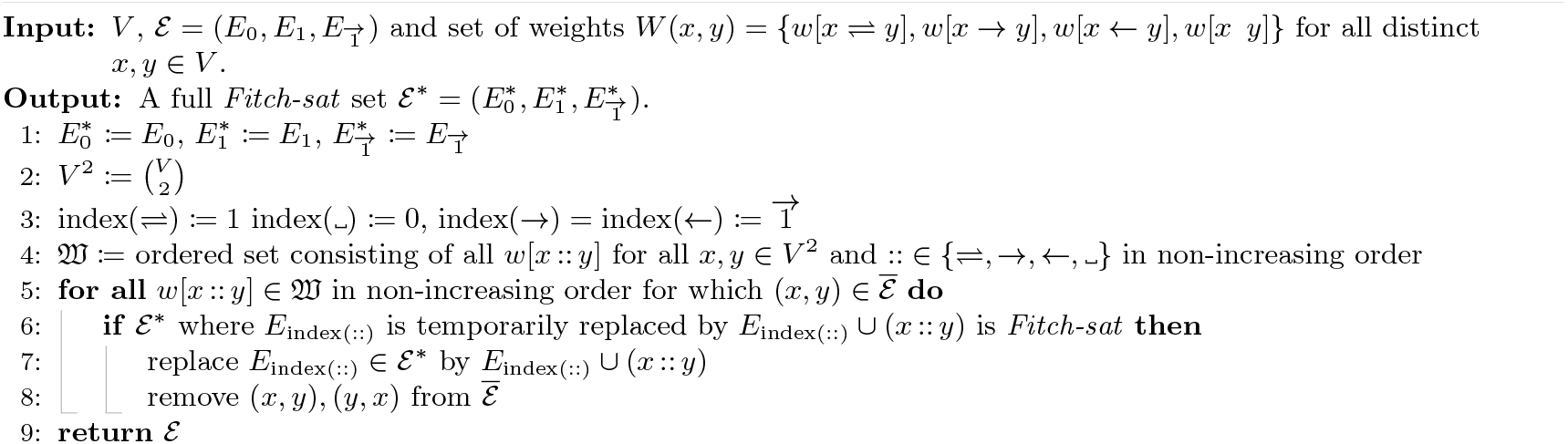

## 6 Computational Results

In this section, we compare the performance of Algorithm 1 and Algorithm 1 in recovering full sets of relations from simulated Fitch graphs. Data and scripts used in this analysis are openly available at https://github.com/bsfaqu/partial-fitch-completion.

Motivated by possible future applications of these algorithms to infer HGT events from sequence data, we generated a synthetic dataset comprising 2100 Fitch graphs (700 each for *n* ∈ {25, 50, 75} vertices). To this end, we simulated gene family histories (GFHs) using the library AsymmeTree [18]. A GFH is represented as a gene tree *T* with *n* leaves embedded into a species tree *S*, see [19, Figure 2]. The tree *S* is simulated using a Yule model (constant speciation rate and null extinction rate) on *n/*2 species. Successively, a gene tree on *n* leaves is built along individual species tree *S* using a constant-rate birth-death process given rates for duplication (D), loss (L), and HGT events (H), where the recipient branch of an HGT event in *S* is chosen randomly. In our simulations, we considered 7 different rates of triples (D, L, H) which are displayed in Figure 5-7. Finally, the tree (*T, λ*) is obtained by removing all branches which lead to losses and by marking all edges *e* whose endpoints map to different lineages in *S* (transfer edges) as horizontal transfer event, i.e., we put *λ*(*e*) = 1. The Fitch graph *G* associated with *T* has vertex set *L*(*T*) and a directed arc (*x, y*) if and only if the path from lca_*T*_ (*x, y*) to *y* contains a transfer edge. We refer to *G* as the true graph when comparing it to an estimated Fitch graph *G*. We generated 100 gene trees for every leaf size and mutation rate combination (*D, L, H*), resulting in the aforementioned 2100 fitch graphs.

For each simulated Fitch graph *G*, we define the true full 3-tuple of relations 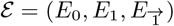 where

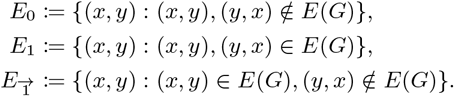

Then, 10%, …, 90% of relations are removed randomly from 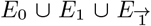. Removing an assignment between vertices *x* and *y* from *E*_0_, resp., *E*_1_ means that both (*x, y*) and (*y, x*) have been removed from *E*_0_, resp., *E*_1_ to maintain the symmetry of *E*_1_ and *E*_0_. Finally, 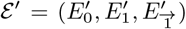 is the partial set of ε containing 90%, …, 10% of the *original information* in ε. This partial tuple ε ^*′*^ serves then as input for Algorithm 1 and Algorithm 1 that are used to obtain an estimated full tuple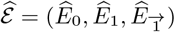. In contrast to Algorithm 1, Algorithm 1 requires a weighting scheme. We used a *constant weighting scheme (* ConstW) and a *“biologically motivated” weighting scheme (* BioW) to evaluate Algorithm 1.

For ConstW, all pairs 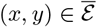 were assigned the same weight *w*(*x* :: *y*) = 0 for all :: ∈ {⇌, →, ←, }. BioW: For *v* ≺_*T*_ *𝓁*, we denote by *𝔈*_*𝓁,v*_ and *ℌ*_*𝓁,v*_ the total number of edges, and the number of transfer edges in (*T, λ*) along the unique path from *𝓁* to *v*, respectively. Moreover, we put (*𝔈* − *ℌ*)_*𝓁,v*_ := *𝔈*_*𝓁,v*_ − *ℌ*_*𝓁,v*_ as the number of edges in *𝔈*_*l,v*_ that are different from transfer edges. Based on this, we define four weights and two *confidence values* that are the basis for the final weights. For all pairs of leaves *x, y* in (*T, λ*), we set

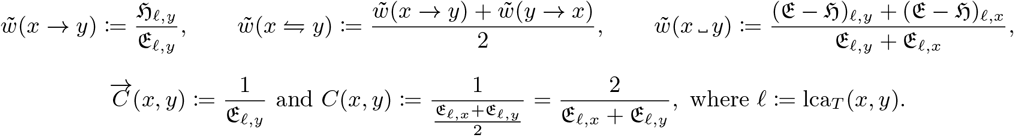

It is straightforward to verify that *w*(*x* → *y*) (resp., *w*(*x* ⇋ *y*)) increases with an increasing number of transfer edges that are along the path from lca_*T*_ (*x, y*) to *y* (resp., both paths from lca_*T*_ (*x, y*) to *y* and to *x*). In contrast, *w*(*x y*) increases with a decreasing number of transfer edges. All of the weights *w*(*x* → *y*), *w*(*x* ⇋ *y*), and *w*(*x y*) are in the real-valued interval [0, 1] and the value 1 is achieved precisely if all edges are transfer edges on the respective paths for the first two weights and if there are no transfer edges for the latter weight. The confidence values 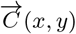 and *C*(*x, y*) are inversely proportional to the respective path length which ensures that the confidence in biologically relevant information regarding the relationship of *x* and *y* is often higher as the divergence time between *x* and *y* becomes smaller. The final weights for BioW that serve as input for Algorithm 1 are obtained by combining the weights and confidence values for all pairs of leaves *x, y* in (*T, λ*) as follows:

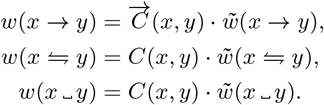

To measure the accuracy of the algorithms, we compute the average difference of the original xenologous pairs in 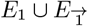 and the estimated ones in 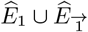 by computing

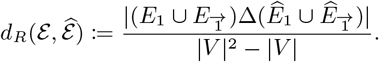

Note that Δ denotes the usual set-theoretic symmetric difference and |*V* |^2^ − |*V* | is the total number of pairs in 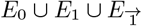, i.e., all elements of *V* × *V* without the reflexive pairs (*x, x*), *x* ∈ *V* .

Figure 3 shows the accuracy, as well as the execution time depending on the percentage of missing information, averaged over all runs of the 700 instances for each *n* ∈ {25, 50, 75}. All the computations were performed on a computer with an Intel(R) Core(TM) i7-6700 CPU @ 3.40GHz (8 cores but not parallelized) and 16GB of RAM. It can be observed that the accuracy results for all three algorithms (Algorithm 1, Algorithm 1 with ConstW and BioW as input) are very similar when no more than 80% (70% for *n* = 25) of information is missing. However, in case of more than 80% (70% for *n* = 25) missing information, Algorithm 1 with BioW outperforms the other algorithms in terms of accuracy. This is possible due to the fact that the weights and confidence values implicitly contain information about the underlying GFH and thus serve as an informative guide to choose the correct pairs of edges added to obtain the estimated Fitch graph. In all cases, the more information of the original Fitch graph is available, the more accurate is its reconstruction.

**Figure 3:**
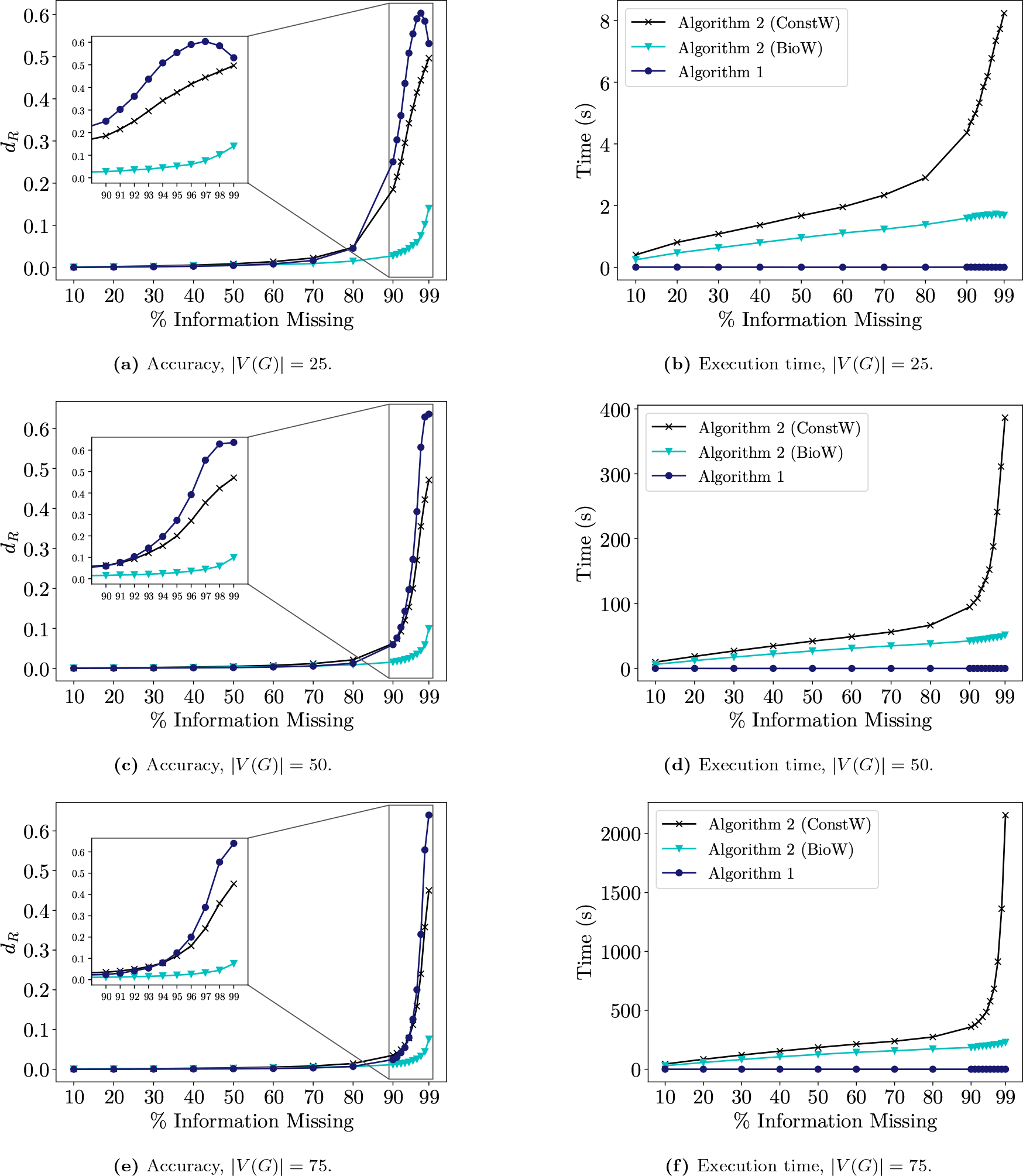
Performance of algorithms over all instances. Shown is the accuracy *d*_*R*_ and execution time depending on the percentage of missing information averaged over all runs of the 700 instances for each *n* ∈ {25, 50, 75}.

Algorithm 1 with ConstW is slower compared to the other two algorithms. In particular, Algorithm 1 outperforms the other algorithms in terms of execution time. Even with an increasing number of vertices, the execution time of Algorithm 1 always stays below one second. In contrast, for Algorithm 1, the running time increases drastically with an increasing number of vertices, which implies that the development of more efficient algorithms taking weights into account is indispensable. The latter observations are also supported by Figures 5, 6 and 7 in the appendix giving a more detailed overview of the influence of different mutation rate combinations (*D, L, H*). Different mutation rates only seem to have a small impact on the performance.

The application of the “rules” (S1), (S2), and (S3) in Algorithm 1 by construction follows a fixed preference order: first (S1), then (S2), and then (S3). According to Lemma 4.1, any applicable rule could be selected in each step. In order to investigate whether this order of the rule application has an influence on the results presented in Figure 3, we modified Algorithm 1 to allow different rule preferences. Figure 4 shows the performance of Algorithm 1 depending on different rule preference orders. The accuracy of the results turn out not to be affected by alternative rule preferences as long as not more than 80% of information is missing. In all cases where more than 80% of the information is missing, the order of rules where (S3) precedes (S2) outperforms all other rules in terms of accuracy. In contrast, when (S2) precedes (S3), then there is a significant decrease of the runtime.

**Figure 4:**
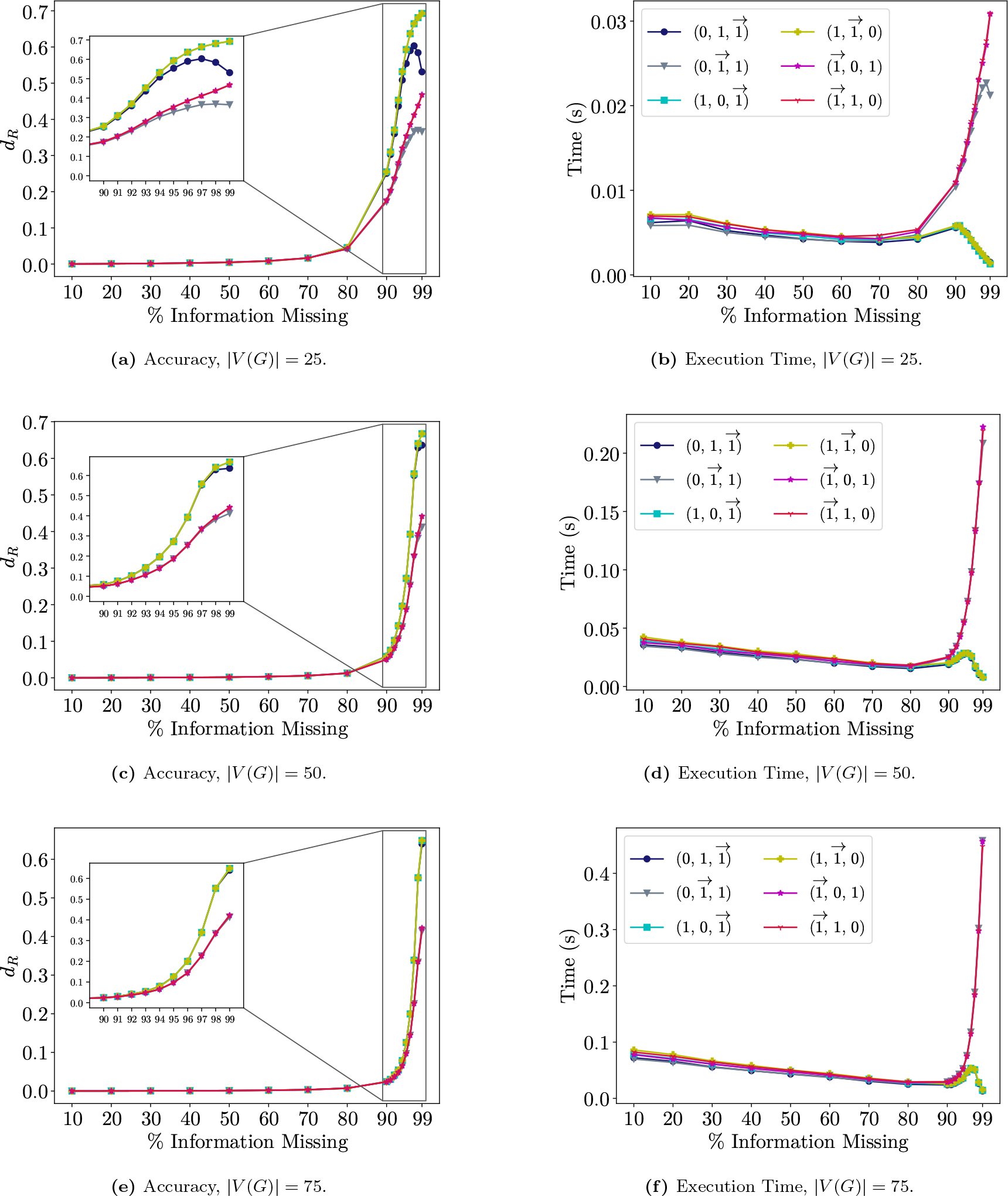
Performance of different rule orders over all instances. Shown is the accuracy *d*_*R*_ and execution time depending on the percentage of missing information averaged over all runs of the 700 instances for each *n* ∈ {25, 50, 75}. The order of rules used in Algorithm 1 is denoted by the preference for the vertex type in the Fitch cotree, i.e., 0 for (S1), 1 for (S2) and 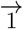 for (S3).

**Figure 5:**
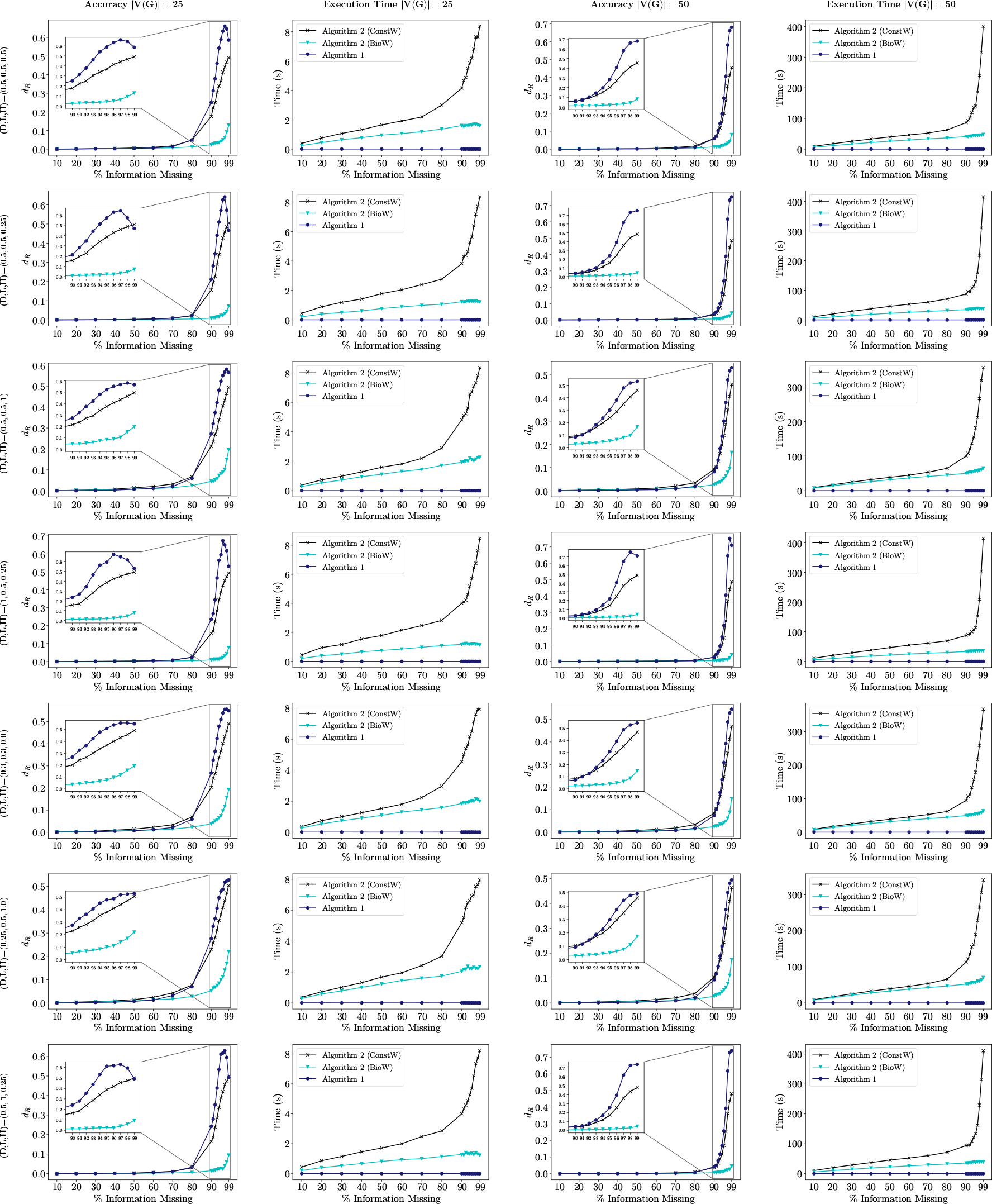
Performance of algorithms over different evolutionary rates for |V(G)| ∈ {25, 50}. Shown is the accuracy *d*_*R*_ and execution time depending on the percentage of missing information averaged over 100 instances for each different evolutionary rate for each *n* ∈ {25, 50}. Evolutionary rates (D,L,H) are chosen for each duplication (D), loss (L), and HGT events (H) from the real-valued interval [0, 1] and are annotated on the left of each row.

**Figure 6:**
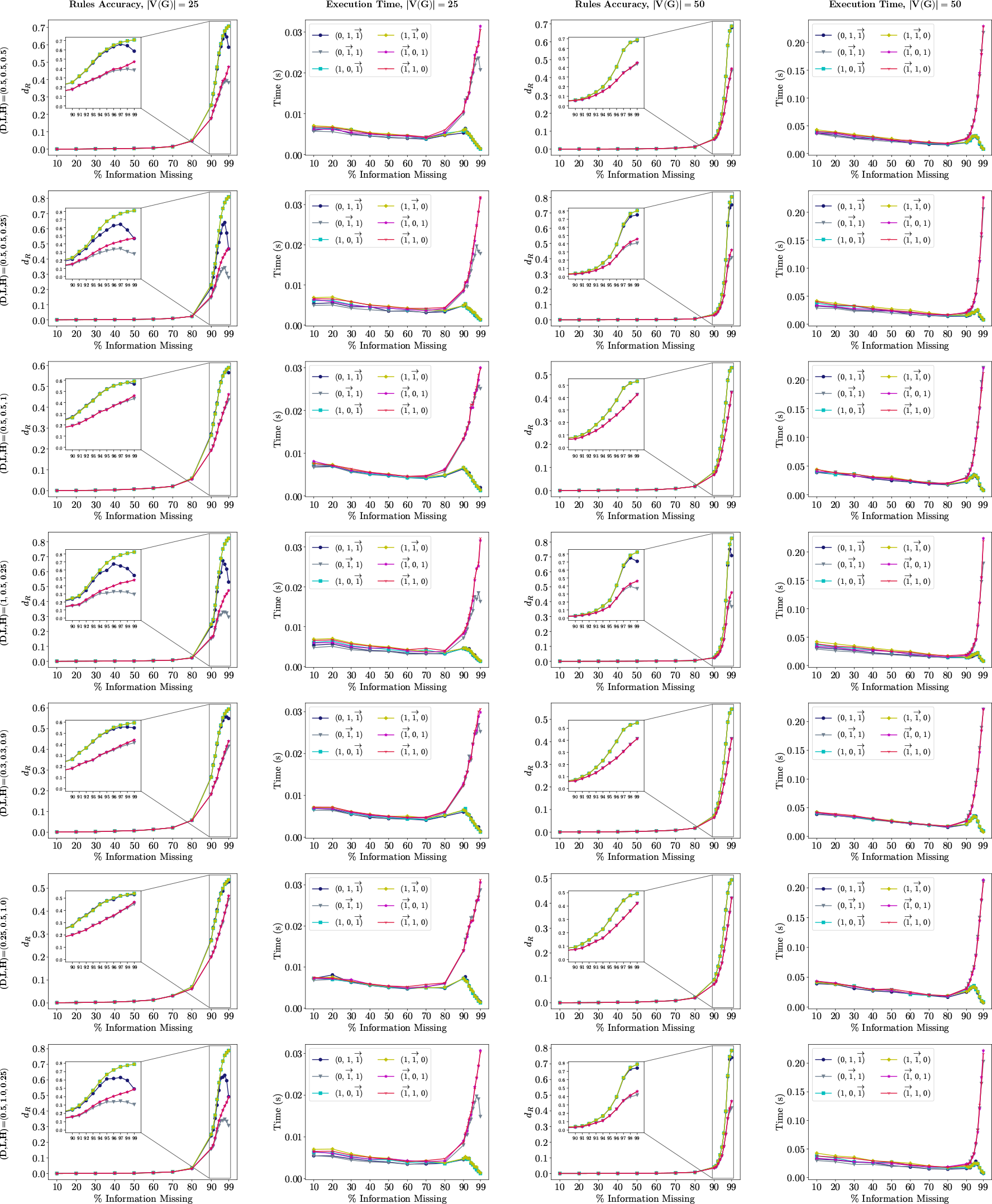
Performance of different rule orders over different evolutionary rates for |V(G)| ∈ {25, 50}. Shown is the accuracy *d*_*R*_ and the execution time depending on the percentage of missing information averaged over 100 instances for each different evolutionary rate for each *n* ∈ {25, 50}. Evolutionary rates (D,L,H) are chosen for each duplication (D), loss (L), and HGT events (H) from the real-valued interval [0, 1] and are annotated on the left of each row. The order of rules used in Algorithm 1 is denote by the preference for the vertex type in the Fitch cotree, i.e., 0 for (S1), 1 for (S2) and 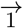 for (S3)

**Figure 7:**
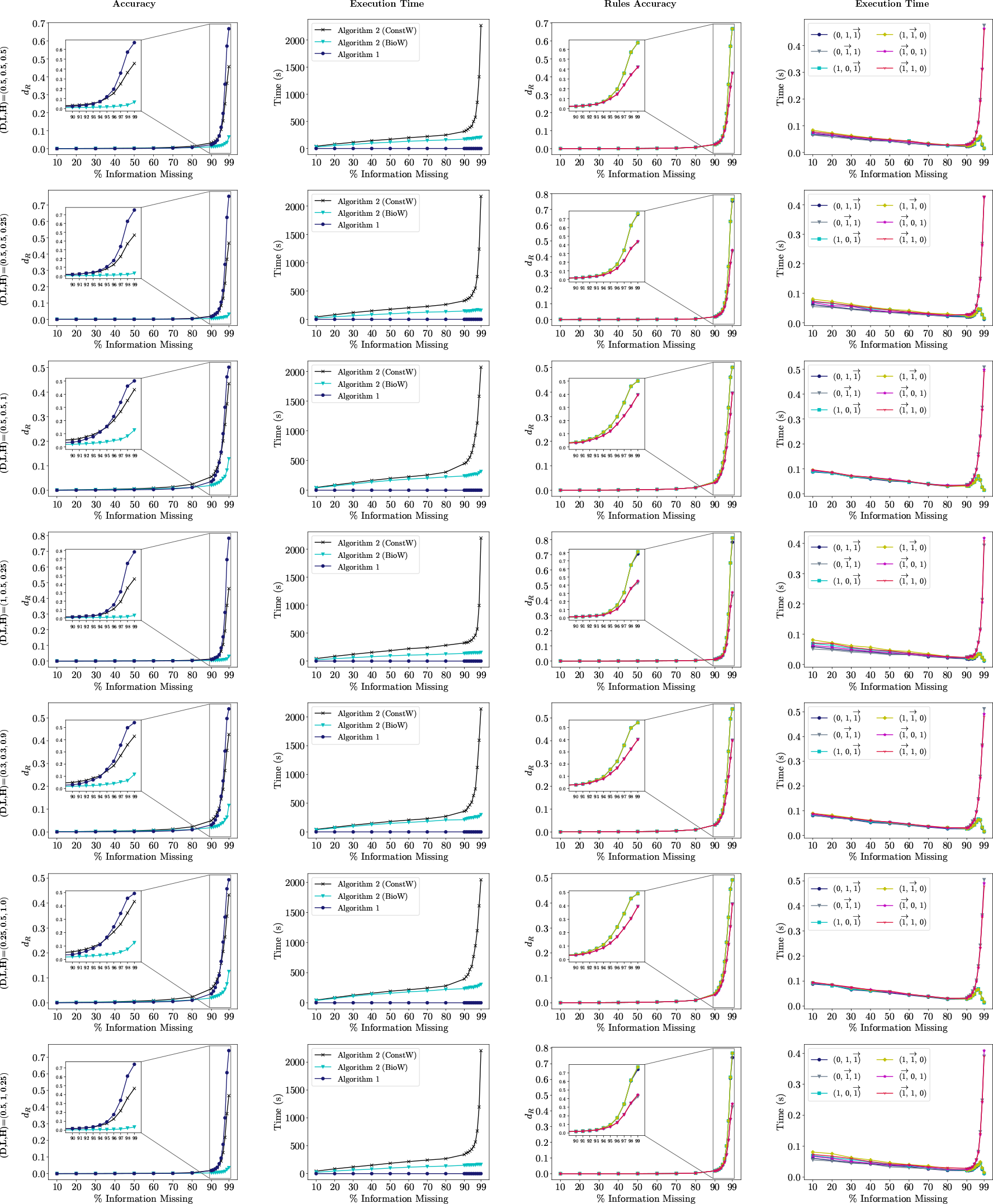
Performance of algorithms and different rule orders over different evolutionary rates for |V(G)| = 75. The accuracy *d*_*R*_ and the execution time, depending on the percentage of missing information, averaged over 100 instances for each different evolutionary rate for *n* = 75. Evolutionary rates (D,L,H) are chosen for each duplication (D), loss (L), and HGT events (H) from the real-valued interval [0, 1] and are written on the left of each row. The order of rules used in Algorithm 1 is denote by the preference for the vertex type in the Fitch cotree, i.e., 0 for (S1), 1 for (S2) and 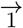 for (S3).

## 7. Concluding Remarks

The characterization of partial Fitch graphs in Theorem 3.11 in terms of Fitch-satisfiable tuples directly leads to the polynomial-time Algorithm 1. In the affirmative case, this algorithm yields a Fitch-cotree, which in turn defines a Fitch graph that contains the given partial Fitch graph as a subgraph. The recognition problem for partial Fitch graphs thus can be solved efficiently. The related Fitch completion problem, which in addition requires optimization of the score function (1), on the other hand is NP-hard (Theorem 5.3). Moreover, we showed that seemingly simpler variants of this problem, such as FC without bi-directional edges (see Theorem 5.5) and FC without non-adjacent vertices, remain NP-complete (see Theorem 5.7).

To tackle these NP-hard problems in practice, we developed a simple greedy heuristic (see Algorithm 1). We evaluated the performance of Algorithm 1 as well as Algorithm 1 on simulated data. Interestingly, we found that only information about a small fraction of correct edges of the original Fitch graph *G* is required to fully recover *G* in most cases. While the accuracy of the greedy algorithm with a biologically inspired weighting schemes is very high, the running time increases drastically with an increasing number of vertices.

The design of more efficient algorithms is thus indispensable for application to large gene families and/or large numbers of genomes.

Moreover, in practice, even a confident empirical estimate of a partial Fitch graph will likely contain errors. In this case, the partial input ε will not be satisfiable. This naturally leads to the question of whether maximal (weighted) Fitch-satisfiable subsets ε ^*′*^ ⊂ ε can be determined efficiently. We suspect this problem, which we will tackle elsewhere, is also computationally hard.

## Acknowledgements

This work was supported in part by the *Deutsche Forschungsgemeinschaft* (grant no: STA 850/49-1).

## A Supplementary Material

In the main text, we included only the performances of the completion algorithms and the effect of the order of the rules on Algorithm 1 averaged across all 7 triples (D,L,H) of evolutionary rates. Here, for each of the evolutionary rates and |*V* (*G*)| ∈ {25, 50}, we report the performances of the completion algorithms in Figure 5 and the effect of the order of the rules on Algorithm 1 in Figure 6. The same results for the case |*V* (*G*)| = 75 are shown in Figure 7. All the computations were performed on a computer with Intel(R) Core(TM) i7-6700 CPU @ 3.40GHz (8 cores but not parallelized) and 16GB of RAM.

